# Photoacoustic Imaging for Non-Invasive Assessment of Physiological Biomarkers of Intestinal Injury in Experimental Necrotizing Enterocolitis

**DOI:** 10.1101/2023.10.20.563296

**Authors:** Jared A. Weis, Jessica L. Rauh, Maryssa A. Ellison, Nildris Cruz-Diaz, Liliya M. Yamaleyeva, Cherrie D. Welch, Kristen A. Zeller, Victoria G. Weis

**Author notes:** Co-corresponding Authors: Jared A. Weis, PhD, 575 N. Patterson Ave, Ste 530, Winston-Salem, NC 27101, Victoria G. Weis, PhD, 391 Technology Way NE, Winston-Salem, NC 27101. These authors contributed equally (Jared A. Weis, Victoria G. Weis).

## Abstract

**Background:** Necrotizing enterocolitis (NEC) is an often-lethal disease of the premature infants’ intestinal tract that is exacerbated by significant difficulties in early and accurate diagnosis. In NEC disease, the intestine often exhibits hypoperfusion and dysmotility, which contributes to advanced disease pathogenesis. However, these physiological features cannot be accurately and quantitively assessed within the current constraints of imaging modalities frequently used in the clinic (plain film X-ray and ultrasound). We have previously demonstrated the ability of photoacoustic imaging (PAI) to non-invasively and quantitively assess intestinal tissue oxygenation and motility in a healthy neonatal rat model. As a first-in-disease application, we evaluated NEC pathogenesis using PAI to assess intestinal health biomarkers in a preclinical neonatal rat experimental model of NEC.

**Methods:** NEC was induced in neonatal rat pups from birth to 4 days old via hypertonic formula feeding, full-body hypoxic stress, and lipopolysaccharide administration to mimic bacterial colonization. Healthy breastfed (BF) controls and NEC rat pups were imaged at 2- and 4-days old. Intestinal tissue oxygen saturation was measured with PAI imaging for oxy- and deoxyhemoglobin levels. To measure intestinal motility, ultrasound and co-registered PAI cine recordings were used to capture intestinal peristalsis motion and contrast agent (indocyanine green) transit within the intestinal lumen. Additionally, both midplane two-dimensional and volumetric three-dimensional imaging acquisitions were assessed for oxygenation and motility.

**Results:** NEC pups showed a significant decrease of intestinal tissue oxygenation as compared to healthy BF controls at both ages (2-days old: 55.90% +/-3.77% vs 44.12% +/-7.18%; 4-days old: 56.13% +/-3.52% vs 38.86% +/-8.33%). Intestinal motility, assessed using a computational intestinal deformation analysis, demonstrated a significant reduction in the intestinal motility index in both early (2-day) and established (4-day) NEC. Extensive NEC damage was confirmed with histology and dysmotility was confirmed by small intestinal transit assay.

**Conclusions:** This study presents PAI as a successful emerging diagnostic imaging modality for both intestinal tissue oxygenation and intestinal motility disease hallmarks in a rat NEC model. PAI presents enormous significance and potential for fundamentally changing current clinical paradigms for detecting and monitoring intestinal pathologies in the premature infant.

## Introduction

Necrotizing enterocolitis (NEC) is a devastating intestinal disease affecting the most fragile premature infants, with mortality rates that persist at 20-40%. With the improvement in premature infant survival rates, the incidence of this disease continues to increase. NEC progresses from intestinal inflammation and epithelial barrier damage to necrosis with resultant perforation that can lead to sepsis-induced multisystem organ failure. The pathologic mechanisms responsible for NEC onset are continuing to be elucidated, but our current understanding suggest this disease pathogenesis likely result in part from systemic stress, such as global hypoxic events, bacterial infection, and compromised intestinal barrier. Ultimately, this early-stage damage can lead to ischemic bowel and gut dysfunction, including dysmotility, that can be challenging to assess and manage in this already fragile population.

NEC presents significant clinical diagnostic difficulties due to the challenging disease and clinical setting. Accurate and conclusive early diagnosis of NEC remains elusive, with limited diagnostic confidence complicating timely and effective interventional efforts to prevent disease progression to urgent surgical removal of necrotic intestine. Advanced non-invasive imaging modalities that may provide increased diagnostic precision for intestinal pathologies (e.g., MRI, CT) present significant challenges in infants, especially in the medically complex NICU patient population, with clinical fragility of these patients necessitating a portable, ‘bedside’ capable imaging modality. In most centers, infants must be transported out of the NICU to these scanners located elsewhere in the hospital, placing the infant at risk of not having the appropriately trained personnel and equipment available in the event of a clinical emergency.

Additionally, radiation exposure from CT is of concern, and preterm infants are unable to undergo prolonged MRI scans in the absence of specialized MRI-compatible incubators as they will become profoundly hypothermic. Other non-invasive imaging methods such as ultrasound (US) have been proposed as a more dynamic and sensitive examination to evaluate the infant intestine. While some US studies have shown promise, overall, studies have not yet fully confirmed its utility or established standard diagnostic criteria in NEC. Specifically, the utility of US has been found to be limited by interpretation difficulties, with poor sensitivity and a false negative rate of up to 40% [1]. These limitations may be due to the use of qualitative features observed with US that can have significant variability between operators rather than defined quantitative biomarkers. Thus, serial plain-film abdominal X-ray (XR) examination remains the clinical standard for diagnosis, despite its limitations. XR represents an isolated snapshot, unable to capture dynamic or functional information about the pictured bowel and thus cannot give any indication of the physiological health of the intestinal tissue. Serial XR offers a crude and slow method of assessing bowel function by allowing clinicians to evaluate the degree to which bowel gas patterns are changing. XR radiographic diagnosis is made upon detection of intestinal wall pneumatosis, with or without pneumoperitoneum [2, 3]. By the time this structural damage is seen on serial XRs, it may have progressed to an advanced stage that necessitates urgent surgical management with associated high morbidity and mortality [2].

Late detection of NEC is associated with significant morbidity and mortality [4]. Conversely, false positive diagnosis leads to unnecessary interventions with prolonged bowel rest and antibiotic use that can lead to poor outcomes in this fragile patient population [5, 6]. NEC therefore remains a significant diagnostic challenge, complicating effective early medical management and prevention of disease progression. As the population of premature infants at the highest risk for NEC continues to rise, a critical need has emerged to develop new innovative and clinically translatable diagnostic methods to enable early interventional opportunities to combat this disease. To address this challenge, in recent work, we introduced the first use of photoacoustic imaging (PAI) to interrogate the infant intestine in a preclinical neonatal rat study [7]. PAI leverages the ‘optoacoustic effect’ to generate an image, whereby near-infrared light is applied to tissue and absorbed by chromophores that then undergo thermoelastic expansion. This results in production of acoustic signal that can be visualized using an ultrasound transducer. Naturally occurring (e.g., hemoglobin, melanin, collagen) and exogenously administered (e.g., indocyanine green) chromophores exhibit spectrally distinct absorption patterns that can be separately visualized and quantified using computational analysis. Of particular interest to NEC, the optical absorption spectra of hemoglobin strongly varies based on oxygenation status, which allows PAI to independently assess oxy- and deoxyhemoglobin, allowing for spatial quantification of tissue oxygen saturation. With a hybrid ‘light in – sound out’ approach to imaging, PAI is capable of more molecularly sensitive and specific detection of physiological features over ultrasound imaging and significantly enhanced imaging penetration depth over optical imaging.

Our recent work validated the use of PAI to quantify important physiological biomarkers of intestinal health in a proof-of-concept study to assess intestinal tissue oxygenation and intestinal motility [7]. PAI is well-suited to address complicated NEC diagnostic features with quantification of intestinal hypoxia and intestinal dysmotility that are known physiological hallmarks of NEC that current clinical diagnostic tools are unable to assess. Additionally, significant prior literature supports the potential promise for these physiological features of intestinal tissue oxygenation and intestinal motility to be used as biophysical biomarkers to detect early intestinal injury and monitor NEC disease. Previous studies have demonstrated a pronounced loss of microvasculature in NEC intestines in both animal NEC models and human disease [8, 9], that leads to intestinal tissue hypoxia. Intestinal hypoxia is also a known major precursor and risk factor for the development of NEC [10, 11]. Intestinal tissue oxygenation has also been previously recognized as a potentially important factor for monitoring intestinal health, with many previous studies attempting to implement near-infrared spectroscopy (NIRS) to measure regional intestinal tissue oxygenation [12, 13, 14, 15] for NEC diagnostics. While NIRS has shown promise in preclinical animal studies [16, 17], translation to the clinic has not been as successful, with clinical studies demonstrating a lack of diagnostic precision, as NIRS is significantly limited in penetration depth with no spatial localization. Previous studies have also established decreased peristalsis as a hallmark of intestinal dysfunction and NEC disease in both human NEC and the NEC experimental animal model [18, 19], with decreases in the enteric nervous system having been shown to contribute to this dysmotility [20, 21]. Impaired intestinal transit due to decreased peristalsis is commonly one of the first clinically observed symptoms of intestinal dysfunction and can further exacerbate intestinal damage and disease [22, 23, 24]. However, effective and accurate quantitative clinical monitoring methods are considerably limited [25, 26, 27], impairing accurate detection and staging of intestinal dysfunction. As an emerging imaging modality, PAI has significant promise to address the current clinical limitations of detection for these important biomarkers of intestinal health and NEC disease.

As a follow-up to our initial proof-of-concept study demonstrating the use of PAI in the neonatal rat intestine [7], in this current work, we evaluated the use of PAI measures of intestinal tissue oxygenation and intestinal motility in an experimental animal model of NEC to establish the utility of non-invasive PAI for potential NEC diagnosis. We hypothesized that PAI can quantitatively characterize changes of both the NEC vascular hallmark of intestinal tissue hypoxia and the NEC intestinal functional hallmark of intestinal dysmotility in an important first-in-disease technology development step towards improving the diagnosis and monitoring of NEC in premature infants.

## Methods

### Animals

Timed-pregnant Sprague-Dawley rats were obtained from Charles River Laboratories (Wilmington, MA, USA). Dams were monitored every 6 hours for birth, beginning 36-hours before anticipated birth. Immediately following birth, pups were divided into healthy breastfed (BF) control or experimental NEC cohorts. Pups of both sexes were used in all studies. For pups in the NEC cohort, newborn rat pups were isolated from the dam within 6 hours of birth and an experimental model of NEC was induced for 4 days via hypertonic formula feeding (four times per day), full-body hypoxic stress (5% O2, three times per day), and orally dosed lipopolysaccharide to mimic bacterial stress (4 μg/g, three times within 30 hours of birth), as previously described[28]. NEC animals were housed in a clinical neonatal incubator repurposed for animal studies, to ensure consistent warmth (32°C air temperature) and humidity (>50%) throughout the study duration, placed within a dedicated isolated room within vivarium space.

BF pups remained with their dams in the same isolated rooms as the NEC pups for the duration of studies. BF pups were removed from Dam only for imaging studies and returned following anesthesia recovery. Use of animals in this study adheres to the protocols approved by the Institutional Animal Care and Use Committee of Wake Forest University.

### Ultrasound and photoacoustic imaging

High frequency anatomical US and PAI images were collected using a Vevo 2100 LAZR High-Resolution Ultrasound and Photoacoustic Imaging System (FUJIFILM VisualSonics, Inc., Toronto, ON, Canada). Anesthesia was induced in rat pups using an anesthesia vaporizer system with inhalation of 2% isoflurane in medical air with airflow at 1 L/min. Following induction, anesthesia was maintained at 1% isoflurane in medical air with airflow at 1 L/min throughout imaging studies. Rat pups were placed supine on the heated imaging platform of the Vevo system with continuous physiological monitoring. Imaging was acquired in the sagittal plane using a LZ-550 transducer, with axial resolution of 44 μm and broadband frequency of 32 MHz - 55 MHz. The Vevo 2100 system utilizes a Nd:YAG laser with an optical parametric oscillator, tunable in the NIR range of 680-970 nm, 20 Hz repetition rate, 5 ns pulse width, 45 +/-5 mJ pulse energy, and peak fluence of <20 mJ/cm^2^. Heated and centrifuged ultrasound gel was used to provide acoustic coupling with the abdomen of rat pups. US and PAI imaging acquisition methods and parameters were held constant for all animals used in this study.

### Photoacoustic imaging of intestinal tissue oxygenation

To measure intestinal tissue oxygenation, similar to our previous study [7], PAI imaging was acquired in ‘Oxy-Hemo’ mode with spectral excitation at 750 nm and 850 nm wavelengths. Two-dimensional (2D) image slices of PAI and co-registered US were acquired both at the abdominal midplane. For acquisition of three-dimensional (3D) volumes, a series of 2D image slices were acquired along a perpendicular axis to the sagittal imaging plane with US transducer location controlled automatically via a motorized controller. Image slices with 0.3 mm image spacing, with a total of 50 images across the abdomen, were used to acquire 3D volumes. Assessment of intestinal tissue oxygenation is based on the calculation of oxygen saturation (sO2) from PAI image acquisition measurements of oxyhemoglobin and deoxyhemoglobin concentrations. sO2 is calculated automatically on the Vevo imaging system as the ratio of oxyhemoglobin to total hemoglobin, using Vevo HemoMeaZure and OxyZated post-processing analysis tools. Using the Vevo Lab analysis software (FUJIFILM VisualSonics, Inc., Toronto, ON, Canada), a region-of-interest (ROI) was manually drawn over the intestinal region within the abdomen for each animal to record measures of intestinal tissue oxygenation. ROIs were designated on both 2D midplane acquisitions and 3D volumetric acquisitions, with mean sO2 measurements within the ROI used to designate intestinal tissue oxygenation in both 2D and 3D analysis. BF and NEC rat pups were imaged at 2- and 4-days old (n=5, each group). To avoid interference from the ‘total body’ hypoxia used to induce NEC disease, all NEC pups were imaged at least 7 hours after the last hypoxia session. From our previous study [7], oxygenation levels in healthy pups equalize based on the inspired O2 levels within 5 mins. Therefore, hypoxia to induce NEC disease does not directly interfere with the quantification of tissue oxygenation in our study design.

### Intestinal motility

To measure intestinal motility, as developed in our previous work [7] and motivated by prior studies [29], the abdomen of each rat pup was imaged using the Vevo system and analyzed for intestinal peristalsis deformation. 1 - 1.5 hours prior to the imaging session, indocyanine green (ICG) (Indocyanine Green for Injection, Diagnostic Green, Farmington Hills, MI, USA) was administered as an intraluminal intestinal contrast agent via oral gavage (7.5 mg/kg, diluted in sterile water). Anatomical US images and co-registered PAI images were acquired using cine acquisitions of ∼30 seconds recordings at 6 Hz frame rate to capture intestinal peristalsis motion and contrast agent transit within the intestinal lumen. ICG contrast enhanced PAI signal intensity was recorded using the Vevo system with single spectral excitation at the ICG peak wavelength of 810 nm. Shown in Supplemental Figure 1, multi-slice sagittal 2D US/PAI cine acquisitions were acquired in 5 locations (2.5 mm spacing) along a perpendicular plane across the abdomen and were used to assess volumetric intestinal motility.

**Figure 1.**
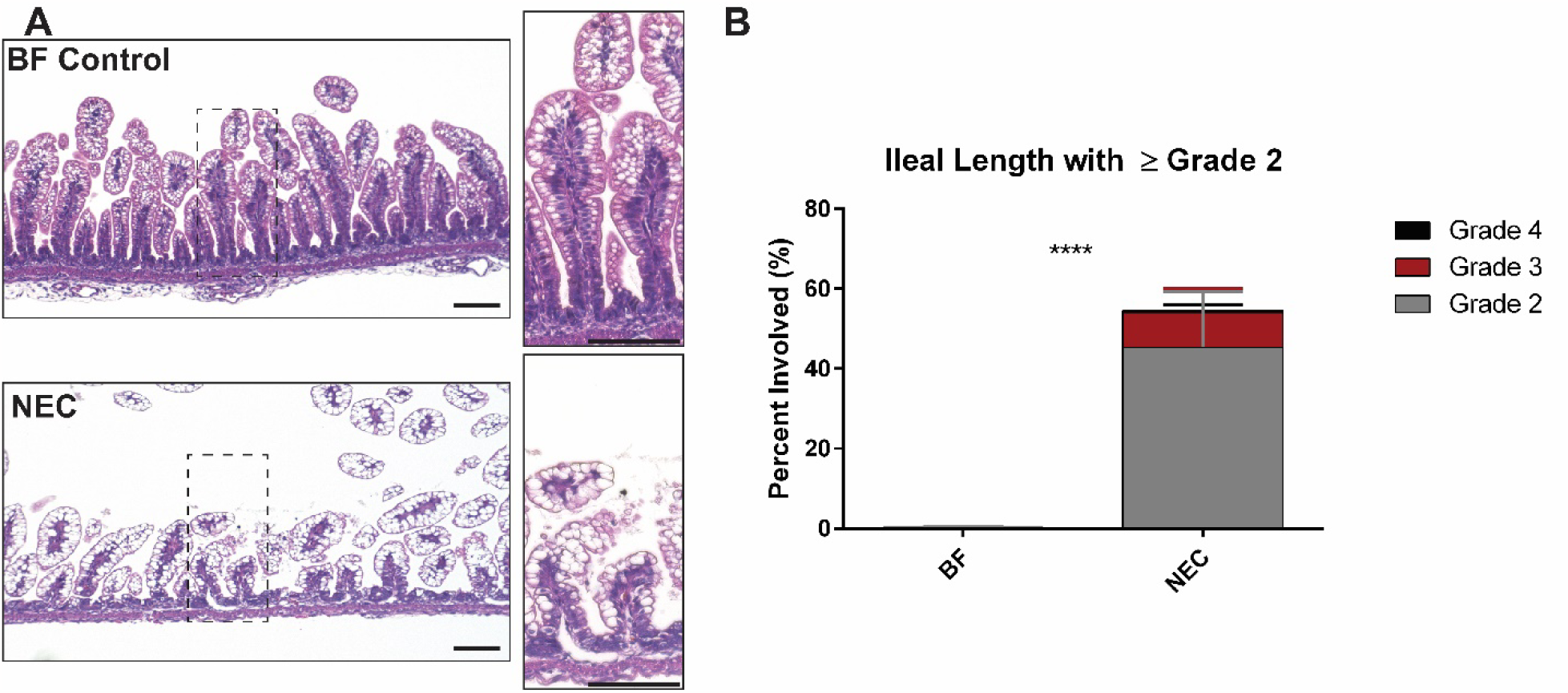
Histological assessment of NEC damage. **A**. H&E of 4-day old rat pup ileum shows that the NEC model induced severe epithelial damage and loss of crypt morphology as compared to BF. Scale bar = 100 μm. **B**. Graph of NEC damage grading along the ileal length shows significant amounts of NEC grade damage (≥ Grade 2) as compared to BF. **** *p*<0.0001.

Intestinal motility was assessed from US and contrast-enhanced PAI images using our previously developed and validated method [7]. Briefly, post-processing image analysis is used to extract motion deformation estimates of intestinal peristalsis throughout the cine time course. Each image in the cine time series is non-rigidly registered to the initial time series image using an initial affine rigid image registration followed by a deformable b-spline based method [30] using the default parameter settings. Deformation is analyzed separately for both US and PAI images. Maps of the intestinal motility index are calculated based on the estimated deformation fields. Deformation estimates at each pixel location are used to construct a Jacobian matrix of the deformation field containing the spatial gradients of the deformation vector.

Expansion/contraction due to intestinal peristalsis is quantified by calculating the determinant of the deformation Jacobian matrix at each pixel within each time frame of the cine time series. Next, the standard deviation calculated in the time dimension for each pixel location is used as a summary measurement of the intestinal motility index. A ROI is manually drawn over the intestinal area within the abdomen based on the anatomical US image and applied to both US and contrast enhanced PAI images. The mean value of the motility index within the intestinal ROI is used as a score of the intestinal motility index. This intestinal motility index is reported at both the single slice abdominal midplane and as an average over all 5 slice locations to designate a volumetric motility index. BF and NEC rat pups were imaged at 2-days old (n=3 and n=7, respectively) and 4-days old (n=3 and n=6, respectively).

### Intestinal transit assay

Following US/PAI imaging data acquisition in both BF and NEC cohorts at 4-days old, intestinal motility was also assessed using a separate gold-standard endpoint physiological assay using a dye-based small intestine-transit assay [31]. Similar to our previous study [7], methylene blue dye (2% aqueous solution, diluted in sterile water) was administered via oral gavage (100 ul volume) to each animal. Dye was allowed to transit for 1 hour, animals were sacrificed via cervical dislocation, and the GI tract from stomach to distal colon was excised and photographed. The distance from the stomach-duodenal junction to the most distal location of methylene blue visibly detected [32] within the 1 hour of transit time was recorded and used as a measure of small intestinal transit velocity.

### Histology

Following all imaging studies in both healthy and NEC animal cohorts at the 4-day old timepoint, intestinal tissues were collected and processed for histological analysis along the entire ileal length, designated as the distal third of the total small intestinal length, as described in our prior study [28]. Hematoxylin and eosin (H&E) staining was performed on 4-μm thick sections containing cross-sections of the entire ileal length and were imaged at 10X using an EVOS FL Auto 2 Cell Imaging System (Invitrogen, CA). H&E stained sections from each animal were graded for intestinal damage according to a histological NEC damage grade severity scoring system, as used in our prior study [28], with 0 designated as normal, 1 as disarrangement of villus cells with mild villus core separation, 2 as disarrangement of villus cells with severe villus core separation, 3 as epithelial sloughing, and 4 as bowel necrosis or perforation. The extent of ileal involvement of each NEC grade was serially assessed along 2-3 cm ileum length segments and the total ileal involvement for each NEC grade was calculated as the total linear length of ileum with each NEC grade, normalized by the total linear length of the ileum. NEC-like damage is associated with a histological score of 2 or greater.

### Statistical analysis

For intestinal tissue oxygenation and intestinal motility metrics at both 2- and 4-days old, the mean value of measurements within the intestinal ROI were compared groupwise between the BF and NEC cohorts using an unpaired t-test. All analyses were completed using GraphPad Prism software with *p*-value < 0.05 considered as statistically significant. For correlation analysis between 2D and 3D imaging analysis metrics, the Pearson’s correlation coefficient was calculated using a pooled analysis to combine all animals from both the BF and NEC cohorts at both 2- and 4-days old.

## Results

### Histology confirms NEC damage

To verify that the experimental animal model of NEC disease induced prominent epithelial damage commonly seen in clinical NEC disease, we examined H&E staining of collected ileal tissues in BF and NEC pups. As previously reported in our recent study [28] and others [33, 34, 35], the experimental NEC model induced aberrant intestinal cell disruption and loss of normal crypt/villus morphology at 4-days old, as shown in Figure 1. NEC-grade damage, with histological NEC score of 2 or greater was found along 55% of the ileal length. In addition to histological evidence of epithelial damage, clinical symptoms consistent with NEC, including dehydration, abdominal distention, diarrhea, and weight loss were observed in NEC pups, similar to our previous studies [28].

### PAI assessment of tissue oxygenation reveals reduced intestinal tissue oxygenation in NEC

We evaluated PAI assessment of intestinal tissue oxygenation in experimental NEC disease as compared to healthy BF control. Representative images for PAI assessment of 2D intestinal tissue oxygenation at the abdominal midplane are shown in Figure 2 for NEC pups as compared to BF control pups at both 2-day (Figure 2A) and 4-days (Figure 2B) old. PAI images qualitatively demonstrate marked reduction in oxygen saturation in NEC vs BF at both 2- and 4-days old. While no visual differences are apparent in the localization and pattern of tissue oxygenation saturation, the magnitude of oxygenation showed large reductions. Post-processing analysis of PAI images yielded quantitative measurements of average tissue oxygenation within the intestinal ROI. NEC pups had a significant decrease of intestinal oxygenation as compared to healthy BF controls at both 2-days (55.90% ± 3.77% vs 44.12% ± 7.18%, p<0.05) and 4-days old (56.13% ± 3.52% vs 38.86% ± 8.33%, p<0.01), reflecting a 21% and 31% reduction in intestinal oxygenation at 2- and 4-days old, respectively.

**Figure 2.**
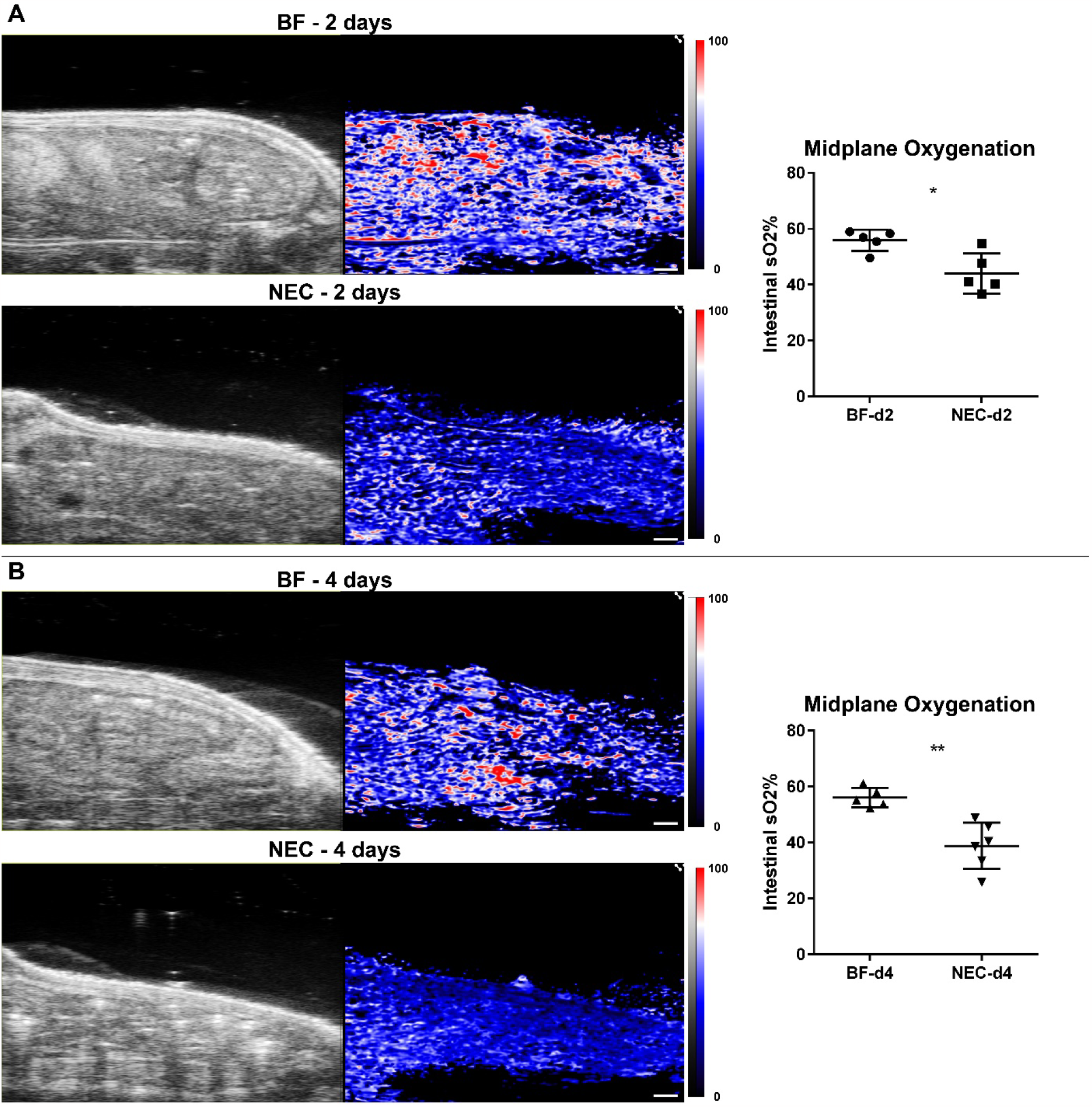
**(Left)** US and PAI of tissue oxygenation (sO2) in representative BF and NEC 2-day (A) and 4-day (B) old rat pups. Scale bar = 1 mm (Scale bar denotes scale for both US and PAI). **(Right)** Intestinal tissue oxygenation in both 2-day (A) and 4-day (B) old NEC pups demonstrated a significant reduction compared to BF controls. \**p*<0.05, \*\**p*<0.01.

To more comprehensively interrogate PAI assessment of intestinal tissue oxygenation in experimental NEC disease, we extended our analysis to full-volumetric 3D analysis. Representative images for PAI assessment of volumetric 3D intestinal tissue oxygenation are shown in Figure 3 for NEC pups as compared to healthy BF control pups at 2- and 4-days old. Visually, marked reduction in volumetric tissue oxygenation is apparent when comparing NEC to BF at both 2- and 4-days old. 3D intestinal ROI quantitative analysis showed that NEC pups exhibit a significant decrease of intestinal oxygenation as compared to healthy BF controls at both 2-days (57.66% ± 2.34% vs 46.81% ± 6.14%, p<0.01) and 4-days old (59.87% ± 3.01% vs 39.37% ± 5.53%, p<0.001), reflecting a 19% and 34% decrease, respectively. These data demonstrate a significant reduction in intestinal tissue oxygenation in both early and established NEC.

**Figure 3.**
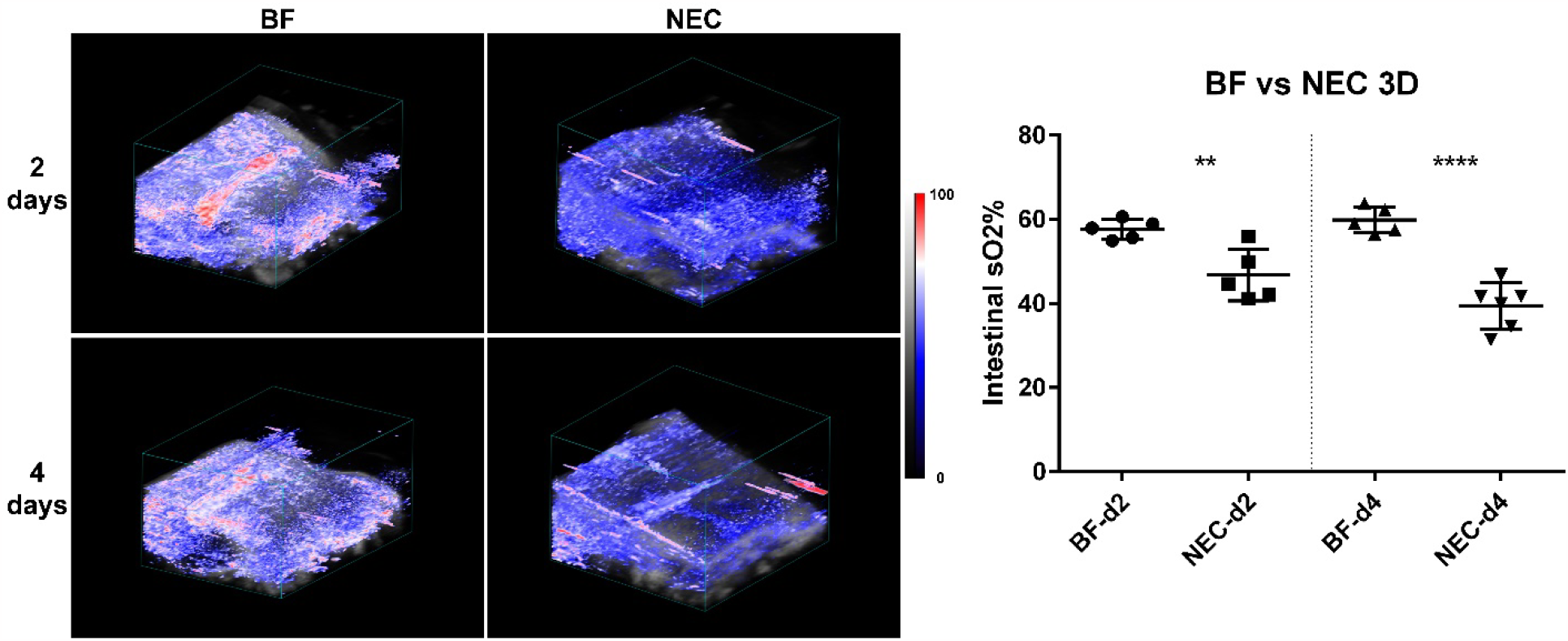
**(Left)** Volumetric assessment of intestinal tissue oxygenation in representative BF and NEC pups at both 2- and 4-days old. **(Right)** Quantification of volumetric intestinal tissue oxygenation demonstrated a significant reduction in NEC as compared to BF control at both 2- and 4-days old. \*\**p*<0.01. \*\*\**p*<0.001.

### US and PAI assessment of peristalsis reveals reduced intestinal motility in NEC

We assessed the intestinal motility index using both US and PAI in NEC animals at 2- and 4-days old, compared to healthy BF controls. To enhance PAI signal intensity of intestinal luminal contents, ICG contrast agent was administered to both cohorts as an intraluminal contrast agent. Videos demonstrating cine US/PAI motility for representative BF and NEC pups at both 2- and 4-days old are shown in Supplementary Videos 1 – 4. As shown in Figure 4, the intestinal motility index was significantly reduced in NEC as compared to BF in 2-day old rat pups as assessed by both US (0.074 a.u. ± 0.019 vs. 0.15 a.u. ± 0.004, p<0.001) and contrast-enhanced PAI (0.11 a.u. ± 0.025 vs. 0.19 a.u. ± 0.002, p<0.001). These motility decreases reflect a reduction of 74% by US assessment and 42% by PAI assessment in the motility index.

**Figure 4.**
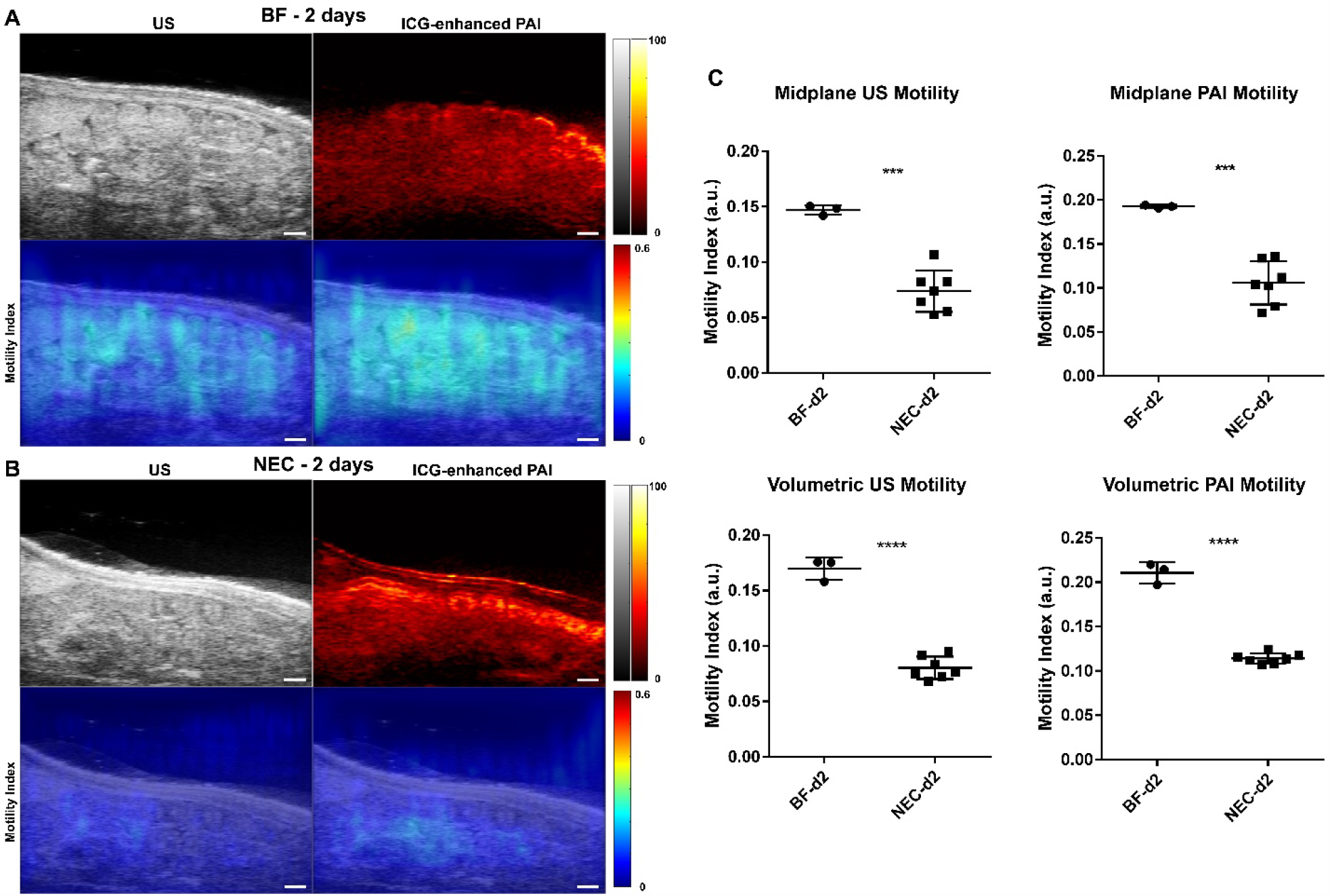
**(Left)** US and ICG-enhanced PAI images (top row) and intestinal motility index maps (bottom row) in representative BF (A) and NEC (B) 2-day old rat pups. Scale bar = 1 mm. **(Right)** Intestinal motility imaging in 2-day old rat pups showed significant reductions in the motility index in NEC animals as compared to BF control. Note that US and PAI motility index measures are to be separately compared due to differences in imaging data source. \*\*\**p*<0.001. \*\*\*\**p*<0.0001.

Similarly, the intestinal motility index was also significantly reduced in NEC compared to BF in 4-day old rat pups assessed using US (0.076 a.u. ± 0.045 vs. 0.16 a.u. ± 0.017, p<0.05) and contrast-enhanced PAI (0.10 a.u. ± 0.042 vs. 0.21 a.u. ± 0.008, p<0.01), as shown in Figure 5.

**Figure 5.**
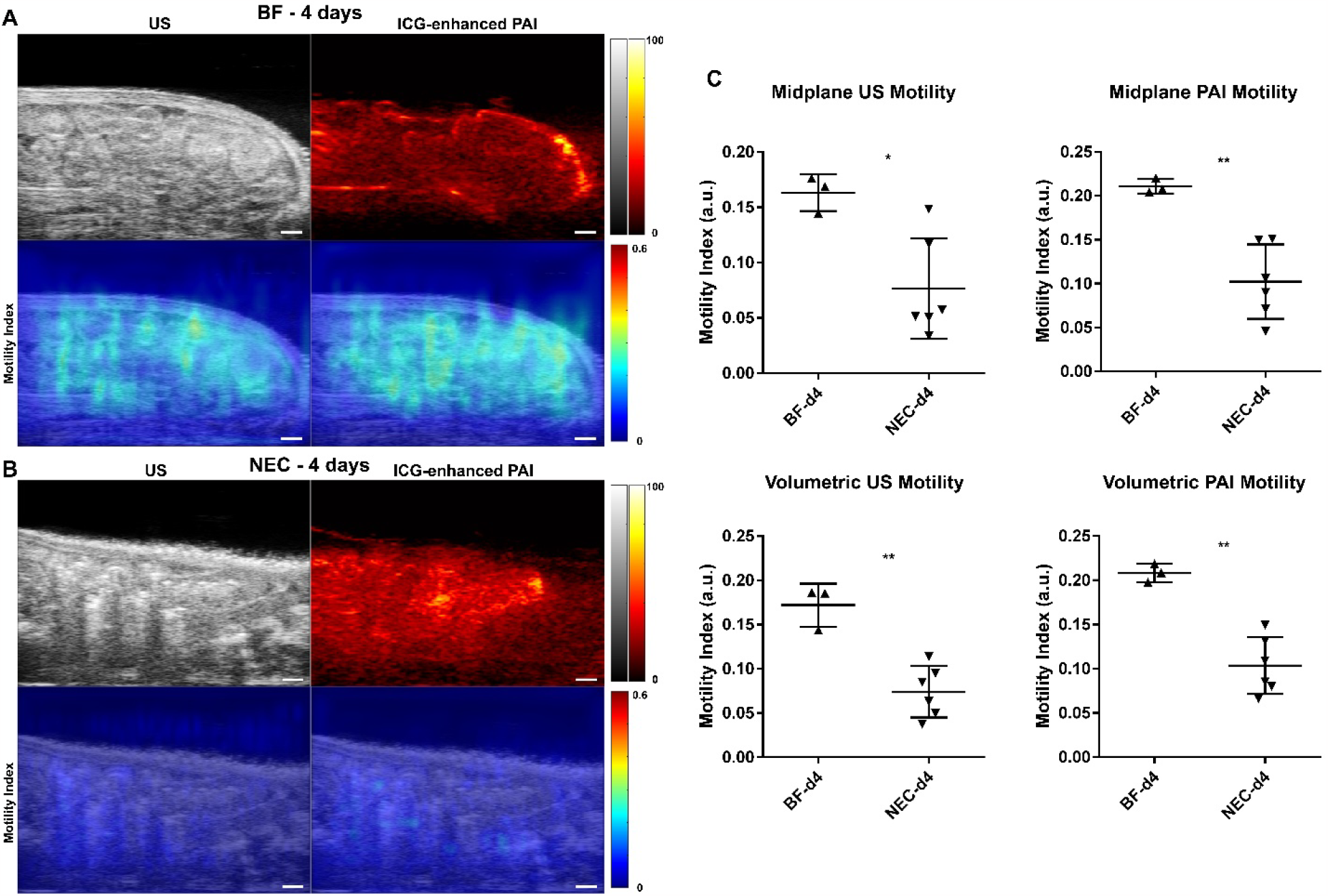
**(Left)** US and ICG-enhanced PAI images (top row) and intestinal motility index maps (bottom row) in representative BF (A) and NEC (B) 4-day old rat pups. Scale bar = 1 mm. **(Right)** Intestinal motility imaging in 4-day old rat pups showed significant reductions in the motility index in NEC animals as compared to BF control. Note that US and PAI motility index measures are to be separately compared due to differences in imaging data source. \**p*<0.05. \*\**p*<0.01.

This is a 53% and 52% decrease in the motility index, respectively. Note that due to differences in the source imaging modality, direct comparisons of the motility index measures between US and PAI are not appropriate.

Assessment of the volumetric intestinal motility index, defined as the average intestinal motility index from five equally spaced single slice acquisitions (as shown in Supplementary Figure 1), showed a similar reduction in the volumetric motility index as found in the 2D midplane motility index. The volumetric intestinal motility index was significantly reduced in NEC compared to BF in 2-day old rat pups, assessed using both US (0.080 a.u. ± 0.010 vs 0.17 a.u. ± 0.010, p < 0.0001) and PAI (0.11 a.u. ± 0.006 vs 0.21 a.u. ± 0.011, p < 0.001) as well as in 4-day old rat pups assess using US (0.074 a.u. ± 0029 vs 0.17 a.u. ± 0.024, p < 0.01) and PAI (0.10 a.u. ± 0.032 vs 0.21 a.u. ± 0.010, p < 0.0001). Reduced intestinal motility in NEC pups as compared to BF was confirmed in 4-day old animals via a methylene blue small intestinal transit assay (Supplemental Figure 2).

### Correlation of 2D and 3D measures of intestinal tissue oxygenation and intestinal motility

We also investigated the correlation between two- and three-dimensional measurements of intestinal oxygenation and intestinal motility using a pooled analysis with all imaged pups, including both BF and NEC cohorts at both 2- and 4-days. Shown in Figure 6A, midplane single slice 2D measures of intestinal oxygenation were found to very strongly correlate with volumetric 3D measures, with a Pearson’s correlation coefficient of 0.94. Correlation analyses to assess the relationships between the midplane 2D and volumetric 3D intestinal motility index for US and PAI are shown in Figure 6B and Figure 6C, respectively. Midplane measurements of the intestinal motility index were very strongly correlated with the volumetric motility index, with Pearson’s correlation coefficients of 0.91 and 0.92 for US and PAI, respectively.

**Figure 6.**
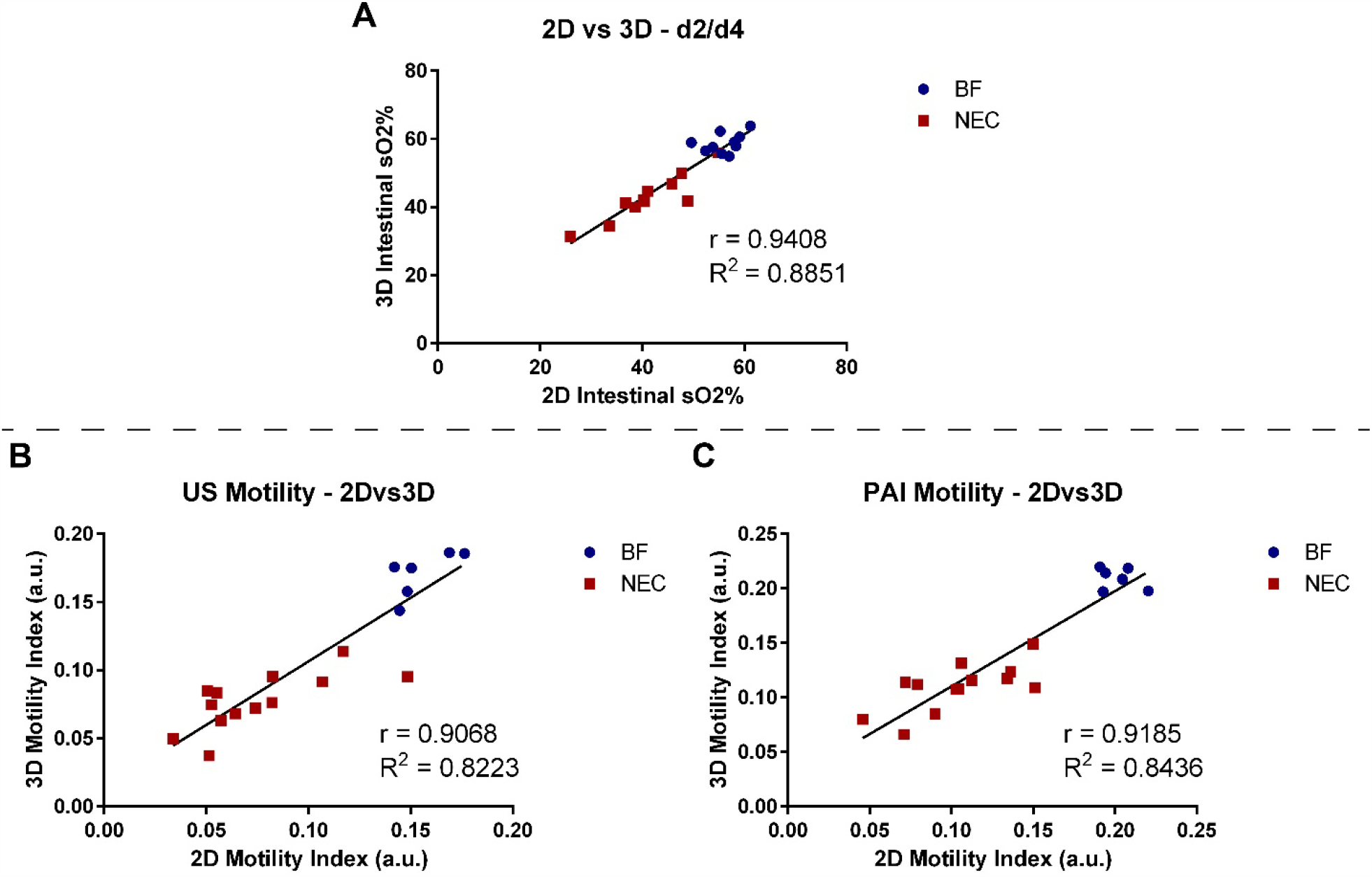
Correlation plots comparing intestinal tissue oxygenation analyzed in 2D vs 3D (A) and correlation plot comparing the intestinal motility index for US (B) and ICG-contrast enhanced PAI (C) analyzed in 2D vs 3D.

## Discussion

The neonatal rat experimental NEC model used in this study recapitulates many of the clinical symptoms and histological damage found in early-stage human NEC disease and has been previously shown to recapitulate major hallmarks seen in human NEC disease [34, 35, 36].

Specifically, this experimental NEC model has been shown to have aberrant microvasculature with increased tissue hypoxia and decreased intestinal motility [8, 9, 10, 20, 22, 23, 24]. With evaluation timepoints sampled at 2-days and 4-days following disease model initiation, this NEC model represents a time course from early disease initiation to established NEC, a timeline for which clinical staging and diagnostic confidence remains limited. Our group recently developed non-invasive imaging and analysis methods that incorporate the novel use of PAI to assess intestinal tissue oxygenation and motility as biomarkers of infant intestine health and disease. In this current study, we now sought to investigate the use of PAI to assess intestinal tissue oxygenation and intestinal motility in our experimental NEC animal model in an important first-in-disease preclinical study. Overall, our NEC animal model recapitulates hallmarks of intestinal dysfunction and human NEC that PAI is well-suited to non-invasively detect and robustly measure. Our studies demonstrate that PAI provides adequate sensitivity and resolution to detect these known changes in intestinal tissue oxygenation and intestinal motility with NEC disease induction and progression. This study demonstrates the feasibility and exceptional promise for the use of PAI to non-invasively assess oxygenation and motility in the healthy and diseased infant intestine, providing key quantification of important biomarkers for intestinal health and disease.

Intestinal tissue hypoxia is both an important risk factor for the onset of NEC disease, as well as a critically important disease mechanism driving subsequent intestinal tissue damage and necrosis [10, 37, 38]. However, current clinical diagnostic imaging methods using XR and US are unable to assess tissue oxygenation, and emerging uses of NIRS are limited by poor penetration depth, lack of spatially resolved measures, and lack of diagnostic precision. We have previously demonstrated the ability of PAI to measure changes in intestinal oxygenation in healthy rat pups with an inspired gas challenge of 5%, 21%, and 100% oxygen. In this current study, we have now assessed tissue oxygenation in NEC disease using PAI. In both 2- and 4-day old rat pups, PAI analysis measured significant reductions in intestinal tissue oxygenation of NEC rat pups as compared to healthy BF controls. This study confirms the ability of PAI to non-invasively and quantitatively detect changes in intestinal tissue oxygenation that are known to occur in NEC disease. While NEC disease can often present both clinically and preclinically with heterogeneous and ‘patchy’ severity within intestinal tissue, we also found significant volumetric reductions in intestinal tissue oxygenation throughout the abdomen in NEC pups, along with very strong correlation of midplane single slice and volumetric measurements of intestinal tissue oxygenation. Overall, these data establish the utility of PAI to quantitively and spatially detect NEC-associated changes in intestinal tissue oxygenation that have not previously been possible via non-invasive methods.

Impaired intestinal transit due to decreased peristalsis, observed as feeding intolerance, is commonly one of the first clinically observed symptoms of intestinal dysfunction[22, 23, 24]. However, effective and accurate quantitative clinical monitoring methods are considerably limited [25, 26, 27], impairing accurate detection and staging of intestinal dysfunction. Using deformation analysis in both US B-mode imaging and intraluminal contrast-enhanced PAI, we quantified significant decreases in the intestinal motility index in NEC as compared to BF in both 2- and 4-day old pups. This reduction of intestinal motility in NEC was confirmed with a small intestinal transit assay, validating the reduced motility measures assessed using non-invasive imaging. We similarly found volumetric reduction in intestinal motility, along with very strong correlation of midplane single slice and volumetric measures of intestinal motility. Together, these data demonstrate the ability to utilize both US and PAI as a new modality to quantitively assess intestinal motility as a potential biomarker for intestinal damage and dysfunction.

Moreover, this new analysis methodology allows for future in-depth investigations into the effects of motility dysfunction on NEC disease onset and progression. While this study is a first-in-disease use of PAI, there are noted limitations of the current study. Although strong statistical significance was achieved in groupwise comparisons between BF and NEC cohorts, the overall sample size and time points in this study were limited. We do show significant changes in intestinal tissue oxygenation and intestinal motility measures alongside histological and physiological confirmation of intestinal damage and impaired intestinal transit. However, future studies with increased sample sizes and more finely resolved evaluation time points throughout the NEC disease time course are needed to better understand the dynamic changes in NEC onset and progression. It will also be important in future studies to perform extensive large-scale paired histological correlation analyses to identify the cellular and molecular origins and mechanisms that underlie PAI observations of reduced intestinal oxygenation and intestinal motility. Additionally, this study was performed in an experimental NEC disease model in rat pups. The strong correlation between 2D midplane and 3D volumetric quantifications in the rat pup model is possibly due to small size of the pup that may impede resolution of regional differences. Future studies within the preterm infant population or larger animal models will allow for further investigations into regional differences in our measures.

Lastly, while prior work has shown that this experimental rat pup model system serves an important role in investigational and discovery research, future studies in a clinical population are ultimately needed to fully examine the utility of PAI for assessment of NEC in the preterm infant population. Overall, the current study provides the foundational work that PAI is a promising imaging modality that detects key physiological indicators of intestinal health in disease. These studies now allow for future studies to address these noted gaps to further develop and investigate PAI for NEC disease.

New advanced non-invasive imaging methods for NEC in the infant intestine present significant opportunities to improve the quality of care for premature infants in the NICU. Due to a lack of tools for assessing intestinal health in the premature infant, current clinical methods rely on a ‘wait-and-see’ approach for early phase cases with suspicion of NEC disease. While there are many ongoing efforts to advance other established imaging modalities such as US, these methods have been limited to mostly qualitatively identifying structural features and are unable to detect more dynamic physiological changes in NEC disease. PAI presents the novel opportunity to develop a new diagnostic imaging technology that can leverage biophysical physiological hallmarks of NEC disease that are known to occur but are not clinically assessable with current diagnostic tools. Development of PAI for NEC disease could have far-reaching implications for fundamentally changing the current diagnostic paradigm for assessing infant intestinal health and disease. With the emergence of potential non-invasive PAI biomarkers capable of rapid and quantitative assessment of changes in intestinal physiology in addition to the current clinical assessments, we may be able to more rapidly guide clinical care in the future based on detecting and monitoring intestinal tissue oxygenation and intestinal motility in the premature infant.

## Acknowledgements

These studies were supported by the Wake Forest Institute for Regenerative Medicine Pilot Award and supported in part by the National Institutes of Health (NIDDK K01DK125633 to VGY, NHLBI R01HL155420 to LMY), American Gastroenterological Association Research Scholar Award in Health Disparities, and Wake Forest University School of Medicine Faculty Start-Up Funds. We acknowledge the Preclinical Ultrasound and Photoacoustic Imaging Core of Wake Forest University School of Medicine supported in part by the Wake Forest Clinical and Translational Science Institute (NIH NCATS UL1TR001420), NIH ORIP/OD-High End Instrumentation Grant S10 OD012330 and the Hypertension and Vascular Research Center.

## Data availability

The datasets generated and/or analyzed within the current study are available from the corresponding authors on reasonable request.

## Contributions

Study conception and design: JW, VW; Data acquisition: JW, VW, NCD, LY, ME; Data analysis and interpretation: JW, VW; Writing and editing the manuscript: All authors.

## Supplementary materials

**Supplemental Figure 1.**
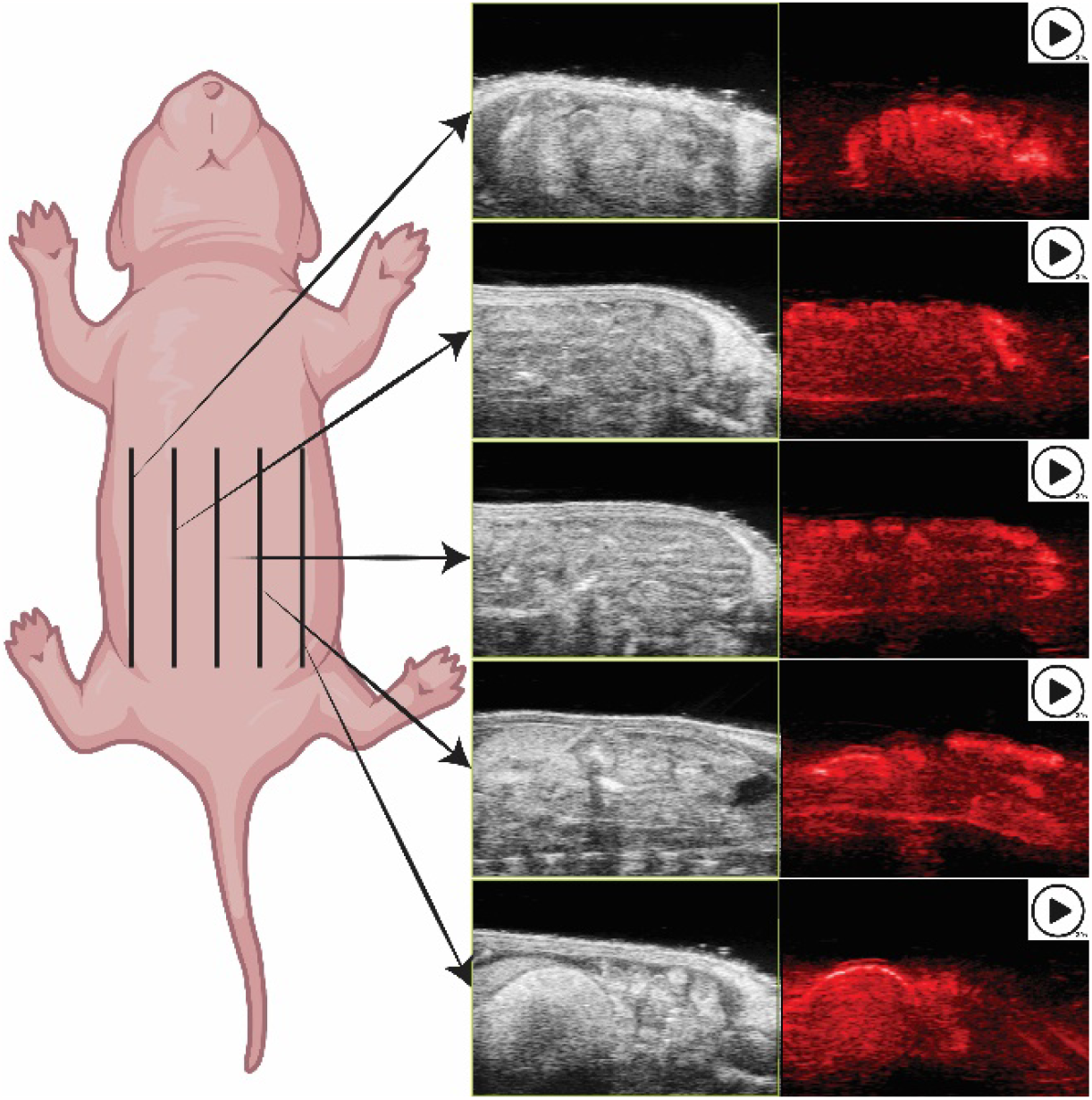
Schematic of volumetric imaging data acquisition for intestinal motility analyses. Volumetric analysis of deformation was performed with 30 second cine US/PAI recordings at 5 equally spaced locations across the abdomen. Lines indicate locations of imaging planes throughout the intestinal volume.

**Supplemental Figure 2.**
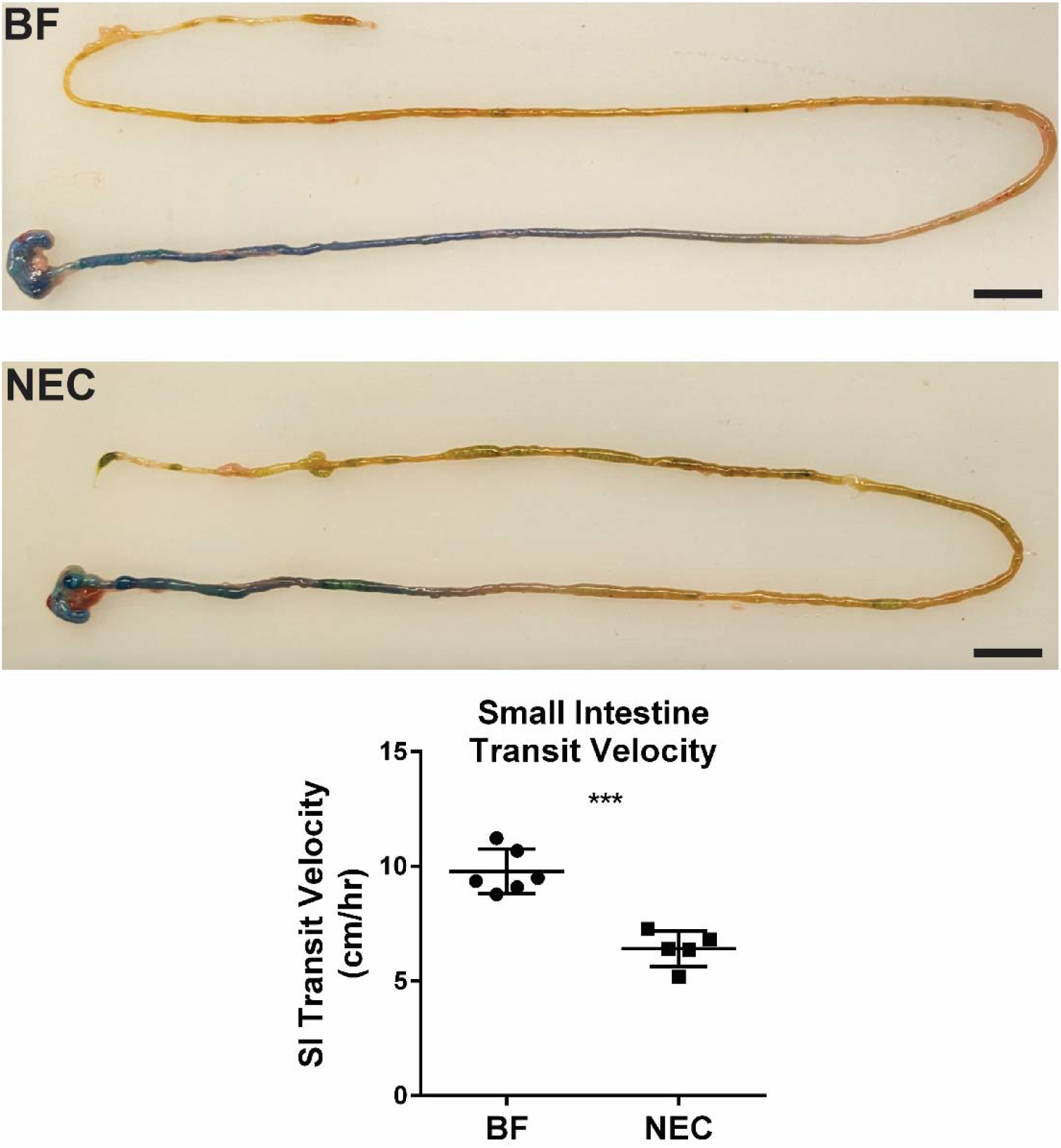
**(Top)** Representative methylene blue images from small intestinal transit velocity assay. Scale bar = 1 cm. **(Bottom)** Transit assay showed a significant reduction in small intestinal transit velocity between BF and NEC. (*p*<0.001). Dye was found to transit significantly less distance in NEC pups as compared to healthy BF pups (6.41 cm ± 0.77 vs. 9.78 ± 0.97, p < 0.001) within the 1-hour transit time.

**Supplemental Video 1**. US and ICG-enhanced PAI imaging for intestinal motility in a representative 2-day old BF pup

**Supplemental Video 2**. US and ICG-enhanced PAI imaging for intestinal motility in a representative 2-day old NEC pup

**Supplemental Video 3**. US and ICG-enhanced PAI imaging for intestinal motility in a representative 4-day old BF pup

**Supplemental Video 4**. US and ICG-enhanced PAI imaging for intestinal motility in a representative 4-day old NEC pup

## References

1 Cuna AC, Lee JC, Robinson AL, Allen NH, Foley JE, Chan SS. Bowel Ultrasound for the Diagnosis of Necrotizing Enterocolitis: A Meta-analysis. Ultrasound Q 2018;34:113–8.

2 Raval MV, Moss RL. Current concepts in the surgical approach to necrotizing enterocolitis. Pathophysiology 2014;21:105–10.

3 Epelman M, Daneman A, Navarro OM, Morag I, Moore AM, Kim JH, et al. Necrotizing enterocolitis: review of state-of-the-art imaging findings with pathologic correlation. Radiographics 2007;27:285–305.

4 Blakely ML, Lally KP, McDonald S, Brown RL, Barnhart DC, Ricketts RR, et al. Postoperative outcomes of extremely low birth-weight infants with necrotizing enterocolitis or isolated intestinal perforation: a prospective cohort study by the NICHD Neonatal Research Network. Ann Surg 2005;241:984–9; discussion 9-94.

5 Stoll BJ, Hansen N, Fanaroff AA, Wright LL, Carlo WA, Ehrenkranz RA, et al. Late-onset sepsis in very low birth weight neonates: the experience of the NICHD Neonatal Research Network. Pediatrics 2002;110:285–91.

6 Cotten CM, Taylor S, Stoll B, Goldberg RN, Hansen NI, Sanchez PJ, et al. Prolonged duration of initial empirical antibiotic treatment is associated with increased rates of necrotizing enterocolitis and death for extremely low birth weight infants. Pediatrics 2009;123:58–66.

7 Weis VG, Cruz-Diaz N, Rauh JL, Ellison MA, Yamaleyeva LM, Welch CD, et al. Photoacoustic Imaging as a Novel Non-Invasive Biomarker to Assess Intestinal Tissue Oxygenation and Motility in Neonatal Rats. Journal of Pediatric Surgery 2023;In press.

8 Ito Y, Doelle SM, Clark JA, Halpern MD, McCuskey RS, Dvorak B. Intestinal microcirculatory dysfunction during the development of experimental necrotizing enterocolitis. Pediatr Res 2007;61:180–4.

9 Senarathna J, Kovler M, Prasad A, Bhargava A, Thakor NV, Sodhi CP, et al. In vivo phenotyping of the microvasculature in necrotizing enterocolitis with multicontrast optical imaging. Microcirculation 2022;29:e12768.

10 Chen Y, Koike Y, Chi L, Ahmed A, Miyake H, Li B, et al. Formula feeding and immature gut microcirculation promote intestinal hypoxia, leading to necrotizing enterocolitis. Dis Model Mech 2019;12.

11 Hackam DJ, Sodhi CP. Bench to bedside - new insights into the pathogenesis of necrotizing enterocolitis. Nat Rev Gastroenterol Hepatol 2022;19:468–79.

12 Fortune PM, Wagstaff M, Petros AJ. Cerebro-splanchnic oxygenation ratio (CSOR) using near infrared spectroscopy may be able to predict splanchnic ischaemia in neonates. Intensive Care Med 2001;27:1401–7.

13 Patel AK, Lazar DA, Burrin DG, Smith EO, Magliaro TJ, Stark AR, et al. Abdominal nearinfrared spectroscopy measurements are lower in preterm infants at risk for necrotizing enterocolitis. Pediatr Crit Care Med 2014;15:735–41.

14 Schat TE, Schurink M, van der Laan ME, Hulscher JB, Hulzebos CV, Bos AF, et al. Near-Infrared Spectroscopy to Predict the Course of Necrotizing Enterocolitis. PLoS One 2016;11:e0154710.

15 van der Heide M, Hulscher JBF, Bos AF, Kooi EMW. Near-infrared spectroscopy as a diagnostic tool for necrotizing enterocolitis in preterm infants. Pediatr Res 2021;90:148–55.

16 Gay AN, Lazar DA, Stoll B, Naik-Mathuria B, Mushin OP, Rodriguez MA, et al. Nearinfrared spectroscopy measurement of abdominal tissue oxygenation is a useful indicator of intestinal blood flow and necrotizing enterocolitis in premature piglets. J Pediatr Surg 2011;46:1034–40.

17 Goldstein SD, Beaulieu RJ, Nino DF, Chun Y, Banerjee A, Sodhi CP, et al. Early detection of necrotizing enterocolitis using broadband optical spectroscopy. J Pediatr Surg 2018;53:1192–6.

18 Halpern MD, Denning PW. The role of intestinal epithelial barrier function in the development of NEC. Tissue Barriers 2015;3:e1000707.

19 Berseth CL. Gut motility and the pathogenesis of necrotizing enterocolitis. Clin Perinatol 1994;21:263–70.

20 Zhou Y, Yang J, Watkins DJ, Boomer LA, Matthews MA, Su Y, et al. Enteric nervous system abnormalities are present in human necrotizing enterocolitis: potential neurotransplantation therapy. Stem Cell Res Ther 2013;4:157.

21 Kovler ML, Gonzalez Salazar AJ, Fulton WB, Lu P, Yamaguchi Y, Zhou Q, et al. Toll-like receptor 4-mediated enteric glia loss is critical for the development of necrotizing enterocolitis. Sci Transl Med 2021;13:eabg3459.

22 Chen W, Sun J, Kappel SS, Gormsen M, Sangild PT, Aunsholt L. Gut transit time, using radiological contrast imaging, to predict early signs of necrotizing enterocolitis. Pediatr Res 2021;89:127–33.

23 Dicken BJ, Sergi C, Rescorla FJ, Breckler F, Sigalet D. Medical management of motility disorders in patients with intestinal failure: a focus on necrotizing enterocolitis, gastroschisis, and intestinal atresia. J Pediatr Surg 2011;46:1618–30.

24 Morriss FH, Jr., Moore M, Gibson T, West MS. Motility of the small intestine in preterm infants who later have necrotizing enterocolitis. J Pediatr 1990;117:S20–3.

25 Gregory KE, Deforge CE, Natale KM, Phillips M, Van Marter LJ. Necrotizing enterocolitis in the premature infant: neonatal nursing assessment, disease pathogenesis, and clinical presentation. Adv Neonatal Care 2011;11:155–64; quiz 65-6.

26 Li YF, Lin HC, Torrazza RM, Parker L, Talaga E, Neu J. Gastric residual evaluation in preterm neonates: a useful monitoring technique or a hindrance? Pediatr Neonatol 2014;55:335–40.

27 Niemarkt HJ, de Meij TG, van de Velde ME, van der Schee MP, van Goudoever JB, Kramer BW, et al. Necrotizing enterocolitis: a clinical review on diagnostic biomarkers and the role of the intestinal microbiota. Inflamm Bowel Dis 2015;21:436–44.

28 Weis VG, Deal AC, Mekkey G, Clouse C, Gaffley M, Whitaker E, et al. Human placental-derived stem cell therapy ameliorates experimental necrotizing enterocolitis. Am J Physiol Gastrointest Liver Physiol 2021;320:G658–G74.

29 Odille F, Menys A, Ahmed A, Punwani S, Taylor SA, Atkinson D. Quantitative assessment of small bowel motility by nonrigid registration of dynamic MR images. Magn Reson Med 2012;68:783–93.

30 Avants BB, Epstein CL, Grossman M, Gee JC. Symmetric diffeomorphic image registration with cross-correlation: evaluating automated labeling of elderly and neurodegenerative brain. Med Image Anal 2008;12:26–41.

31 Welch MG, Margolis KG, Li Z, Gershon MD. Oxytocin regulates gastrointestinal motility, inflammation, macromolecular permeability, and mucosal maintenance in mice. Am J Physiol Gastrointest Liver Physiol 2014;307:G848–62.

32 Miller MS, Galligan JJ, Burks TF. Accurate measurement of intestinal transit in the rat. J Pharmacol Methods 1981;6:211–7.

33 Ares GJ, McElroy SJ, Hunter CJ. The science and necessity of using animal models in the study of necrotizing enterocolitis. Semin Pediatr Surg 2018;27:29–33.

34 Lopez CM, Sampah MES, Duess JW, Ishiyama A, Ahmad R, Sodhi CP, et al. Models of necrotizing enterocolitis. Semin Perinatol 2023;47:151695.

35 Won MM, Mladenov GD, Raymond SL, Khan FA, Radulescu A. What animal model should I use to study necrotizing enterocolitis? Semin Pediatr Surg 2023;32:151313.

36 Sulistyo A, Rahman A, Biouss G, Antounians L, Zani A. Animal models of necrotizing enterocolitis: review of the literature and state of the art. Innov Surg Sci 2018;3:87–92.

37 Zheng L, Kelly CJ, Colgan SP. Physiologic hypoxia and oxygen homeostasis in the healthy intestine. A Review in the Theme: Cellular Responses to Hypoxia. Am J Physiol Cell Physiol 2015;309:C350–60.

38 Singhal R, Shah YM. Oxygen battle in the gut: Hypoxia and hypoxia-inducible factors in metabolic and inflammatory responses in the intestine. J Biol Chem 2020;295:10493–505.

